# Changes in arthropod community but not plant quality benefit a specialist herbivore on plant under reduced water availability

**DOI:** 10.1101/2020.05.25.115519

**Authors:** Po-An Lin, Chia-Ming Liu, Jia-Ang Ou, Cheng-Han Sun, Wen-Po Chuang, Chuan-Kai Ho, Natsuko Kinoshita, Gary W. Felton

**Affiliations:** Department of Entomology, Pennsylvania State University, State College, PA; Graduate School of Life and Environmental Sciences, University of Tsukuba, Ibaraki, Japan; Institute of Ecology and Evolutionary Biology, National Taiwan University, Taipei, Taiwan; Department of Agronomy, National Taiwan University, Taipei, Taiwan; Department of Agro-Bioresources Science and Technology, Faculty of Life and Environmental Sciences, University of Tsukuba, Ibaraki, Japan

**Author notes:** these authors contributed equally to the work (Po-An Lin, Chia-Ming Liu, Jia-Ang Ou). We show that low water availability benefits certain insects through reducing negative species interactions, and highlight the importance of community factors in plant-insect-environment interactions.

**Keywords:** drought, environmental heterogeneity, predation, competition, insect herbivore

## Abstract

Plants grow under reduced water availability can have divergent effects on insect herbivores, in some instances producing benefits to them. However, the forces mediating these positive impacts remain mostly unclear. We conducted a manipulative field study using a specialist herbivore *Pieris rapae*, and its host plant, *Rorippa indica*, in two populations to identify how water availability impacts overall plant quality and multitrophic interactions. We observed that *R. indica* growing under low water availability led to higher survival of *P. rapae* larvae. The increase in survival of eggs and larvae was related to the reduced abundance of other herbivores and natural enemies. Water availability had differential impacts on members of the herbivore community through changes in plant quality. Low water availability decreased the quality of *R. indica* to most herbivores as indicated by reduced abundance in the field and decreased relative growth rate in feeding assays. In contrast, the performance of *P. rapae* larvae were not affected by differences in sympatric *R. indica* grown under different water availability. These results indicate that local *P. rapae* possess some physiological adaptation to overcome fluctuations in host quality. Our findings illustrate that reduced water availability is beneficial to a specialist herbivore, but detrimental to most other herbivores. Our work highlights the complex roles of the arthropod communities associated with plants in determining the impacts of water availability on insect herbivores.

## Introduction

Water availability is a vital factor influencing individuals, populations, and communities of plants (Bunker and Carson 2005; Schulze 1986) and ultimately heterotrophic organisms across trophic levels (Anderegg et al. 2013). Its importance in affecting ecological interactions will likely increase due to the predicted increase in frequency and severity of drought under current models of climate change (Li et al. 2009). In addition to large-scale climatic factors, fine-scale topographic heterogeneity also influences water availability to individual plants, even when the overall climatic conditions are similar (Murren et al. 2020). This is particularly the case for plants in urban and agricultural environments, because conditions such as the reduction in tree canopy coverage, increased in light intensity/temperature, and shallow soil depth all contribute to high evaporation and low water availability (Dambros et al. 2013; Pickett et al. 2011). Given the many dimensions that influence plant water access, the variation in plant phenotype due to changes in water availability is likely a key factor influencing the dynamics and survival of associated insect herbivores.

The influence of water availability on insect herbivores has receive much attention in past literature. A classic example is the relationship between drought and outbreak of insect herbivores (Mattson and Haack 1987; White 1974; White 1969). This observation has led to a fruitful area of research and it is now clear that the impact of water availability on insect herbivores and plants is context-dependent and varies across different systems (Jamieson et al. 2012; White 2009). However, the main factors that contribute to this positive effect of low water availability on some insect herbivores remain unclear. A major limitation in past literature that contribute to this uncertainty was the emphasis on the direct interaction between plants and insects (English-Loeb et al. 1997). In contrast to a beneficial impact observed in natural conditions, low water availability generally elicits negative effects on plant quality (defined as a performance response of insect herbivores, see (Awmack and Leather 2002)) especially in herbaceous plants (Waring and Cobb 1992). It is likely that factors conttributing to these beneficial impacts are interactions apart from the direct interaction between the focal herbivore and its host plant. Low water availability has been documented to reduce the abundance and composition of arthropods associated with plants (Trotter et al. 2008), and can potentially alter the strength and direction of multispecies interactions. Changes in arthropod species interactions (e.g., competition, predation) could affect the fitness of insect herbivores, and have been shown to contribute more to the overall impacts of environmental changes than the effect of pairwise interactions between plants and insect herbivores (Ockendon et al. 2014). Although important, few studies investigating the influence of multitrophic interactions on insect herbivores feeding on plant with different water availability have been performed (Jamieson et al. 2012).

In this study, we aimed to evaluate how water availability alters the performance of a focal insect species through direct effects on plant quality and indirect effects mediated by changes in other arthropods. Our principal hypothesis was that although decreased water availability would reduce the quality of herbaceous plants to insect herbivores, decline in natural enemies and potential competitors could be a factor that positively influences insect herbivores. To test our hypothesis, we conducted a manipulative field study using a wild host, *Rorippa indica* (L.) Hiern, of the specialist herbivore, cabbage white butterfly (*Pieris rapae*) from two populations in Taiwan and Japan. *R. indica* is a perennial glabrous herb that is commonly found in highly disturbed areas such as roadsides, gardens, and field margins throughout the main island of Taiwan (Huang 1996) and also during May to September in most parts of Japan (Jisaburo□ O□i 1965). *R. indica* individuals with different phenotypes due to water availability are commonly observed in the field. By manipulating water status of *R. indica* in the field, in combination with feeding assays in the lab, we aimed to determine (1) the responses (e.g., preference and performance) of insect herbivores to *R. indica* of different water status, and (2) the factors (e.g., plant quality and species interactions) that contribute to these responses.

## Materials and methods

### Insect and plant

Cabbage white butterflies, *Pieris rapae*, were collected in the experimental field of National Taiwan University (NTU, Taipei, Taiwan) and University of Tsukuba (UT, Ibaraki, Japan), and kept at 14:10 light:dark cycle, 25-27°*C*, on cabbage (*Brassica oleracea* var. capitata) and Komatsuna (*B. rapa* var. Perviridis) respectively. North American *P. rapae* were collected on campus at the Pennsylvania State University (PSU, University Park, PA, USA) and maintained on kale (*B. oleracea* var. sabellica). Cabbage looper, *Trichloplusia ni*, corn earworm, *Helicoverpa zea*, and fall armyworm, *Spodoptera frugiperda* were purchased from Benzon Research (Carlisle, PA, USA) and reared on artificial diet (Peiffer and Felton 2005). All insects were kept under the same condition described above. Seeds of *Rorippa indica* were collected in the experimental fields. Plants were maintained in semi-closed greenhouse and grown in 3-inch pots (400 mL) with a mixture of peat moss (Kekkila, Finland): perlite (2-8 mm) (3:1 volume) supplied with fertilizer Osmocote Plus (Everris, Netherland). The water holding capacity of peat moss was estimated as 195 mL (65% of volume) based on information provided by Kekkila.

### Water treatment and water status quantification

Plants with 7-8 fully expanded leaves were subjected to two levels of water treatments: (1) high water availability (200 mL day^−1^ pot^−1^), and (2) low water availability (10 mL day^−1^ pot^−1^). The water levels were chosen based on morphological comparisons of plants from a series of water availabilities (10, 30, 50, 100, and 200 mL day^−1^ pot^−1^) to plants in field growing under moist and dry area. Water was applied daily at 9 a.m. for 7 days. To quantify the water status of plants (Jones 2007), we measured shoot biomass, root biomass, shoot water content, and volumetric water content of soil (*θ_v_*). For shoot biomass and water content, the entire above ground tissue was collected, weighed, dry (70 °C, 48 h), and weighed again. For root biomass, below ground tissue was collected after the above ground tissue, roots were washed and dried (70 °C, 48 h), and weighed to obtain dry mass. For soil water content, 100 cm^3^ of soil sample was collected from each pot avoiding root tissues. Soil was weighed and dried (70 °C, 48 h), and weighed again. The water content was calculated using the following formula: (mass of water_(g)_density of soil_(gcm^−3^)_) (mass of soil_(g)_density of water_(gcm^−3^)_)^1^. All samples were collected 3 h after daily water application. For field experiments, water status of *R. indica* was maintained by the same procedure, accompanied with daily observations at 12, 2, and 4 p.m. to provide more water if the low water availability treatments exhibited low turgor. When signs of low turgor were observed, 10 mL of water was added. This procedure prevented most leaves from irreversible tissue damage. Loss of turgor was not observed for plants growing under high water availability. The phenotypes were maintained as consistent as possible throughout the field experiment. (Appendix 1).

### Oviposition preference, survival, and relative growth rate of insect herbivores

Individual *R. indica* grown under different water status (in pots) were placed in the experimental field at 7 a.m., 3 m apart in a square grid (Appendix 2). Ovipositional preferences were determined by recording the number of eggs of *P. rapae* after the first 72 h. Individual plants were kept in the field for an additional 7 days before *P. rapae* larvae were collected. Only larvae older than second instar were collected to ensure that the collected caterpillars came from the recorded eggs. Because *P. rapae* larvae rarely move away from their host even when the host plant is stressed (based on our rearing observations) and high levels of egg predation, the survival of *P. rapae* was defined and calculated as: (caterpillar number) (egg number)^−1^. See “Method: Arthropod community” for additional details on field experiments.

We analyzed the relative growth rate (RGR) of larvae to determine the quality of *R. indica* to caterpillars under different water treatments. Larvae of *P. rapae* from Taiwan and Japan were fed with *R. indica* from sympatric population. Third instar larvae of *P. rapae* were weighed and placed in 30 c.c. plastic cups with thin layer of 1% agar (5 mL). The larvae were fed with detached mature leaves of specific water status (either 200 mL day^−1^, 10 mL day^−1^, and an additional intermediate water level: 50 mL day^−1^). Leaves were replaced daily to minimize the potential changes in quality caused by excision. Larvae were weighed after 3 days. The start and end weights were used to calculate RGR using the equation: (end weight – start weight) (end weight + start weight)^−1^ days^−1^ (Felton et al. 1989).

We conducted another set of common garden experiment to test the changes in host quality using insects without previous evolutionary history with *R. indica* from Taiwan and Japan. Plants were grown in a greenhouse in PSU with 16:8 light cycle at 25°*C*. Leaves were collected from two different water treatments (200 mL day^−1^, 10 mL day^−1^) from both populations of *R. indica* and fed to third instar larvae of caterpillars from North America, including *P. rapae* (specialist), *Trichloplusia ni* (semi-specialist, cabbage looper), *Helicoverpa zea* (generalist, corn earworm), and *Spodoptera frugiperda* (generalist, fall armyworm) using method as described above.

### Arthropod community

The insect community associated with *R. indica* was determined on the same day by visual observation. All arthropods on each plant during 7 a.m.-12 p.m. were recorded. All morphospecies observed in the field were listed in Appendix 3. Field experiments took place in Taiwan during April – June 2018, and in Japan during July – September 2018. The experiment was replicated three times in both locations, with one trial in each month, defined as early, middle, and late season. The experimental fields were located within the campus of NTU (25°00’55.9”N 121°32’25.0”E) and UT (36°06’51.7”N 140°05’59.7”E), and both were at an elevation of about 900 m^2^. The natural community of plants in the experimental fields was allowed to grow undisturbed. Each trial contained 20-25 individual *R. indica* from each water treatment (total of 40-50 plants). Timing of field experiments were determined according to governmental weather forecasts in Taiwan and Japan. The experiments were conducted during a period of 10 days that has the lowest probability of rain in each season. The incidences of raining were rare, on average 0-1 days per field experiment. In case of rain, plants were covered with transparent plastic bins and removed after the rain. In addition, all plants were placed on a round plastic tray to prevent moisture from the soil. To determine parasitism incidence, larvae of *P. rapae* collected from the field were reared individually using *Brassica oleracea* var. capitata and *Brassica rapa* var. Perviridis as food in Taiwan and Japan respectively until pupation. If the caterpillar died before pupation, the carcass was dissected to determine presence of parasitoids.

### Statistical analysis

All statistical analyses were conducted in R version 3.5.3 (R Core Team 2017). Univariate response variables were modeled with generalized linear models (GLM) with treatment, population, season and their interaction as fixed effects. We analyzed all count data (egg, caterpillar, herbivore, and natural enemy) with negative binomial GLM (package: *MASS*) (Venables and Ripley 2013) with biomass of plants as an offset variable. Survival probabilities of *P. rapae* were modeled with binomial GLM as the odds ratio between alive (caterpillar count) and dead (egg count - caterpillar count). We excluded samples with no eggs observed. Dispersion, zero-inflation, residual normality, and homoscedasticity of all model fits were evaluated with simulation-based residual plots (package: *DHARMa*) (Hartig 2017). Significance of model parameters were tested by Type II rather than Type III Wald’s Chi-squared test (package: *car*) (Fox and Weisberg 2018) to account for unbalanced data (Langsrud 2003).

To examine whether the survival of *P. rapae* could be attributed to abundance of other arthropods, we pooled samples between treatments and fitted binomial generalized linear mixed-effect models (GLMM, package: *glmmTMB*) (Brooks et al. 2017) with observation-level random effect to improve residuals. Abundances of trophic guilds (herbivore and natural enemy, Fig. 4), and the abundances of individual species (Table 2) were used as fixed effect predictors. Individual GLMM was fitted for each season in each location. We implemented backwards model selection (likelihood ratio tests) until all terms were significant (Zuur et al. 2009). Other model diagnostics are performed as described above. Continuous variables such as water content, biomass, and relative growth rate were analyzed using ANOVA, followed by Tukey HSD. Diagnostic plots were used to check for model assumptions, such as equal variance and normality (Zuur et al. 2010). Indicator species analysis was conducted to ascertain whether specific species were associated with a particular water treatment in a particular season (packages: *indicspecies* and *vegan*) (Cáceres and Legendre 2009; Oksanen et al. 2010). We conducted indicator species analysis on presence absence data to partially remove the effects of the difference in absolute abundance among species as we were more interested in whether each individual species showed any association towards a particular treatment-season combination. Permutation test (with 999 permutations) were used to assess the significance of associations between species and treatment-season combinations. In addition, we applied PCA on log(x+1)-transformed (improve skew of species abundance distributions) and z-standardized (put equal weights on each species) species abundances independently for herbivores and natural enemies to help visualize the changes in species composition between treatments and seasons.

## Results

### Water status quantification

Different water treatments led to observable changes in plant phenotype (please see Fig. 1a). Leaf number and shape were not affected by water availability (Fig. 1b); however, leaf shape differed between populations (Fig. 1c, F_1,29_ = 13.03, *P* = 0.001). Fresh weight of shoot of individuals was reduced under low water availability (Fig. 1d, F_1,29_ = 65.29, *P* < 0.001) and higher in the Taiwanese population compared to the Japanese population (F_1,29_ = 22.59, *P* < 0.001). Dry weight was also reduced under low water availability (Fig. 1e, F_1,29_ = 17.05, *P* < 0.001) and higher in the Taiwanese population (F_1,29_ = 8.5, *P* = 0.007). Water content of shoot was reduced under low water availability (Fig. 1f, F_1,28_ = 105.02, *P* < 0.001), and higher in Taiwanese population (F_1,28_ = 5.48, *P* = 0.027). However, the effect of population was only significantly reduced under low water availability, indicated by a significant interaction between water and population (F_1,28_ = 5.48, *P* = 0.027). The biomass of root was not affected by water treatments. However, root biomass was higher in Taiwanese population (Fig. 1g, F_1,28_ = 8.605, *P* = 0.007). Root: shoot ratio was increased under low water availability (Fig. 1h, F_1,28_ = 88.17, *P* < 0.001). Volumetric water content (*θ_v_*) of the soil was 0.53-0.55 for high water availability and 0.09 for low water availability (Fig. 1i, F_1,29_ = 2238.65, *P* < 0.001).

**Fig 1.**
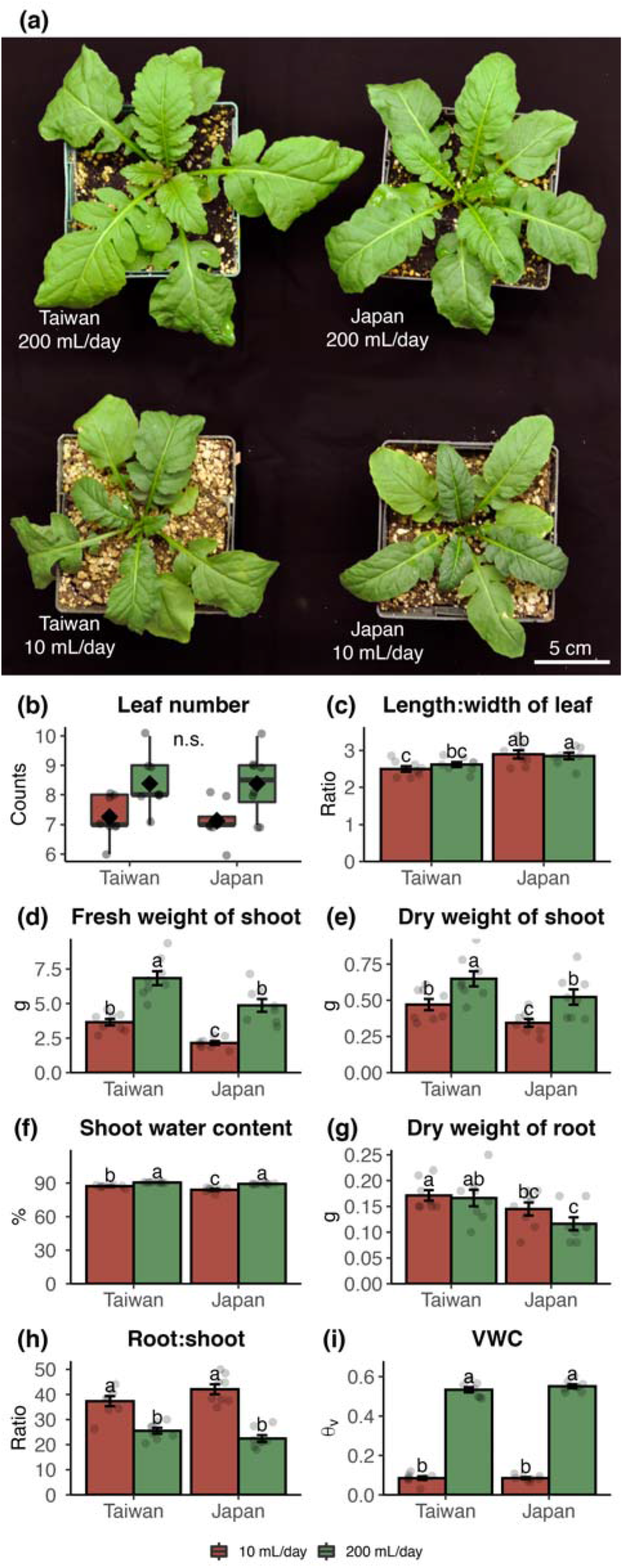
Quantification of water status on *Rorippa indica*. **(a)** phenotypes after water treatment in a common garden (2 h after daily water application). (**b**) Box plot of leaf number (N=8, GLM, Poisson). (**c**) Ratio of length and width of mature leaves (N=8). (**d**) Fresh weight of shoot (N=8). (**e**) Dry weight of shoot (N=8). (**f**) Shoot water content (N=8). (**g**) Dry mass of root (N=8). (**h**) Root: shoot ratio (g g^−1^, dry mass) (N=8). (**i**) Soil volumetric water content (θ_v_) (N=8). Dots indicate individual observations. Values indicate mean ± SE. Different letters indicate significant differences between means (ANOVA, Tukey HSD).

### Oviposition preference, caterpillar number, and survival

Higher quantities of eggs were observed on *R. indica* under high water availability (Fig. 2a, GLM, water: 200 mL day^−1^, estimate = 1.04, Z = 4.79, *P* < 0.001). Egg number on *R. indica* was higher in Taiwan than in Japan (GLM, region: Taiwan, estimate = 2.2, Z= 10.21, *P* < 0.001). In addition, egg number was lower during late season in Taiwan (GLM, season*region: late*Taiwan, estimate = −1.74, Z= −7.11, *P* < 0.001).

**Fig 2.**
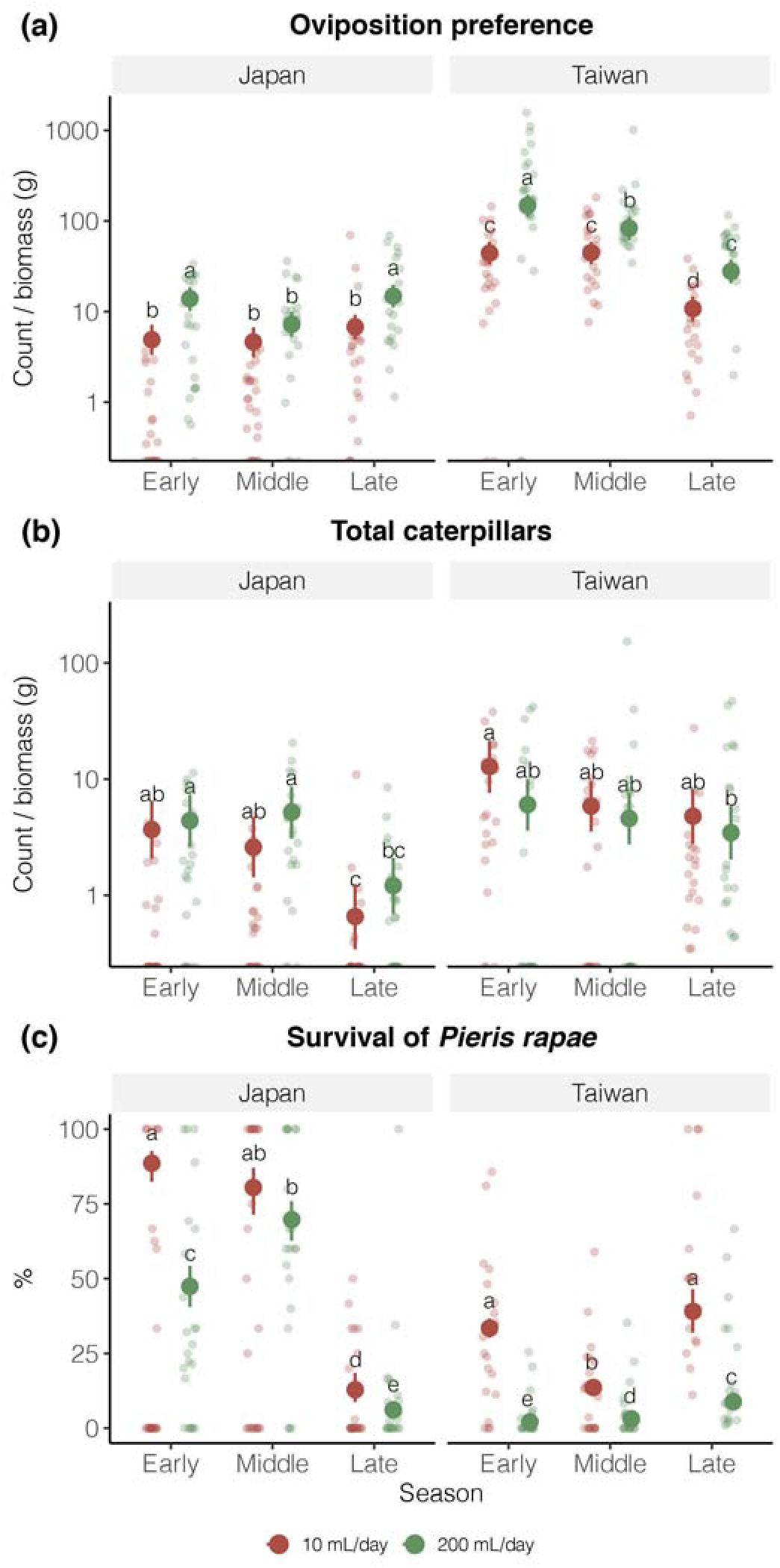
Influence of water status (of *Rorippa indica*), population, and season on *Pieris rapae*. (a) Ovipositional preferences. (b) Total caterpillars at the end of field experiment, (c) Survival of *P. rapae* on *R. indica*. Count data were offset by biomass of *R. indica*. Due to higher values of egg, caterpillar, and herbivore count, data were plotted at the log scale. Transparent points indicate individual observations; opaque points and error bars indicate mean ± 95% CI.

The number of caterpillars on *R. indica* was overall higher in Taiwan (Fig. 2b, GLM, region: Taiwan, estimate = 1.25, Z= 3.57, *P* < 0.001), but lower on *R. indica* under high water availability in Taiwan (GLM, water*region: 20 mL day^−1^*Taiwan, estimate = −0.94, Z= −2.66, *P* = 0.008). The number of caterpillars were lower during late season (GLM, season: late, estimate = −1.72, Z = −4.19, *P* < 0.001), and not affected by water treatment in general.

Survival of *P. rapae* was lower on *R. indica* under high water availability (Fig. 2c, GLM, water: 200 mL day^−1^, estimate = −2.15, Z = −8.79, *P* < 0.001), middle, and late seasons (GLM, season: middle, estimate = −0.63, Z = −2.58, *P* = 0.01; season: late, estimate = −3.96, Z = −14.95, *P* < 0.001). However, the pattern was influenced by several interacting factors. Survival of *P. rapae* was higher on *R. indica* under high water availability in middle and late season (GLM, water*season: 200 mL day^−1^*middle, estimate = 1.57, Z = 10.1, *P* < 0.001; water*season: 200 mL day^−1^*late, estimate = 1.31, Z = 6.52, *P* < 0.001), but lower on *R. indica* under high water availability in Taiwan (GLM, water*region: 200 mL day^−1^*late, estimate = −0.53, Z = −2.34, *P* = 0.019). The survival of *P. rapae* was overall lower in Taiwan (GLM, location: Taiwan, estimate = −2.74, Z = −10.81, *P* < 0.001), but significantly higher in late season (GLM, season*location: late*Taiwan, estimate = 4.2, Z = 17.49, *P* < 0.001) and lower in middle season (GLM, season*location: middle*Taiwan, estimate = −0.53, Z = −2.34, *P* = 0.019).

### Relative growth rate of caterpillars

The relative growth rates (RGR) of *P. rapae* larvae were not affected by water status of *R. indica* from the same geographic origin (Fig. 3a, b). In contrast, populations of *R. indica* from Taiwan and Japan growing under low water availability led to significant decreases in RGR of *P. rapae* from North America (Fig. 3c, F_1_,_141_ = 12.96, *P* < 0.001). *R. indica* growing under low water availability decreased RGR of generalist caterpillars. RGR of *H. zea* larvae were lower on *R. indica* growing under low water availability (Fig. 3d, F_1,147_ = 25.41, *P* < 0.001). RGR of *T. ni* larvae were lower on *R. indica* growing under low water availability (Fig. 3e, F_1,128_ = 8.76, *P* = 0.004) and *R. indica* from Taiwan (F_1_,_128_ = 8.62, *P* = 0.004). However, the negative impacts of low water availability were only significant when *T. ni* larvae fed on *R. indica* from Taiwan (F_1_,_128_ = 5.2, *P* = 0.024). *S. frugiperda* completely rejected *R. indica* from Taiwan as food source, and we observed a small amount of feeding on *R. indica* from Japan. The mortality of *S. frugiperda* larvae were significantly lower when *R. indica* from Taiwan was provided as food (Fig. 3f, GLM, 200 mL day^−1^: estimate = −1.81, *P* < 0.001).

**Fig 3.**
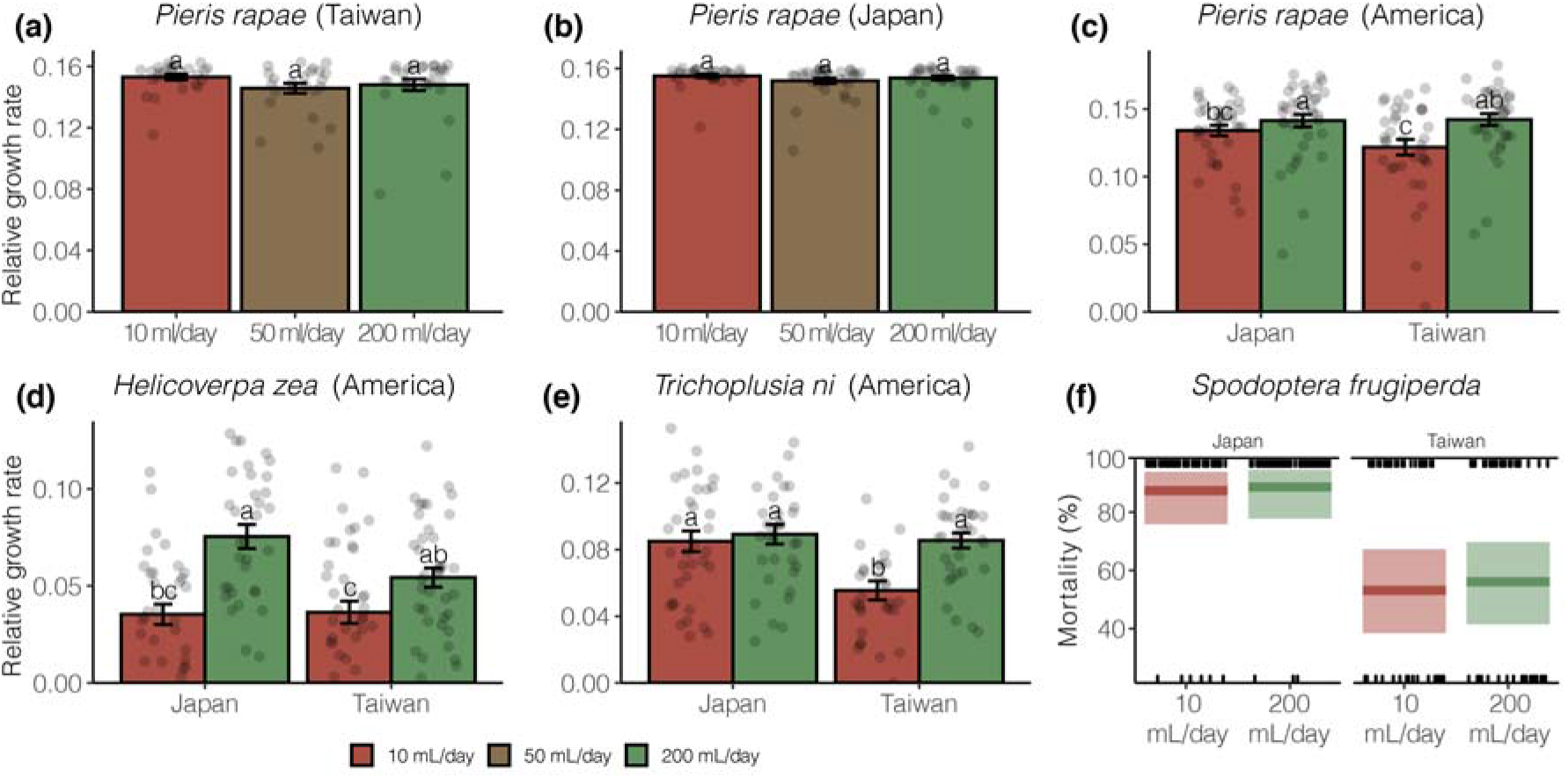
Relative growth rates of lepidopteran larvae feeding on leaves of *Rorippa indica*. (a) *Pieris rapae* (Taiwan) and *R. indica* (Taiwan) (N=30). (b) *P. rapae* (Japan) and *R. indica* (Japan) (N=35). (c) *P. rapae* (North America) and *R. indica* from both location (Taiwan and Japan) (N=40). (d) *Helicoverpa zea* (North America) and *R. indica* from both location (Taiwan and Japan) (N=40). (e) *Trichoplusia ni* (North America) and *R. indica* from both location (Taiwan and Japan) (N=40). Values represent means ± SE. Different letters indicate significant differences between means (ANOVA, Tukey HSD). (f) *Spodoptera frugiperda* (North America) and *R. indica* from both location (Taiwan and Japan) (N=40). 1 = dead, 0 = lived. Bar indicated the mean and boxes represent 95% CI (GLM, binomial).

### Abundance and composition of herbivores and natural enemies

Herbivore abundance (Fig. 4a and Fig. 4i) on *R. indica* was higher on *R. indica* growing under high water availability (GLM, water: 200 mL day^−1^, estimate = 1.43, Z = 4.32, *P* < 0.001), and lower in late season (GLM, season: late, estimate = −2.61, Z = −6.54, *P* < 0.001). The negative association between herbivore abundance and late season was especially stronger on *R. indica* growing under high water availability (GLM, water*season: 200 mL day^−1^*late, estimate = −1.45, Z = −3.39, *P* < 0.001), and became positive during late season in Taiwan (GLM, season*location: late*Taiwan, estimate = 2.69, Z = 6.23, *P* < 0.001).

**Fig 4.**
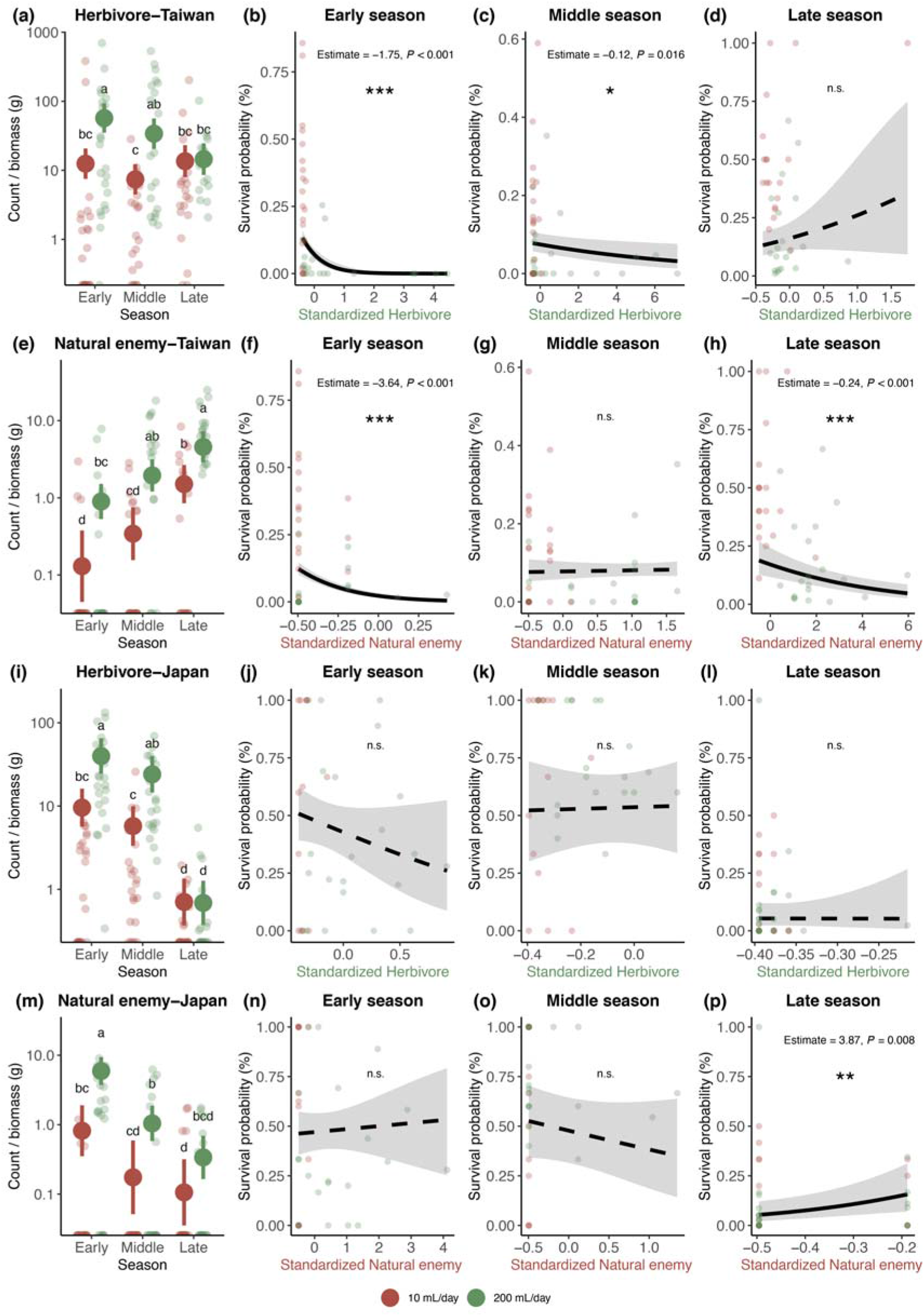
Abundance of herbivores and natural enemies and its associations with water status of *Rorippa indica* and survival of *Pieris rapae*. (a) Abundance of herbivore in Taiwan. (b-d) Association between herbivore abundance and survival of *Pieris rapae* in Taiwan during early, middle, and late season. (e) Abundance of natural enemy in Taiwan. (f-h) Association between natural enemy abundance and survival of *Pieris rapae* in Taiwan during early, middle, and late season. (a) Abundance of herbivore in Japan. (b-d) Association between herbivore abundance and survival of *Pieris rapae* in Japan during early, middle, and late season. (e) Abundance of natural enemy in Japan. (f-h) Association between natural enemy abundance and survival of *Pieris rapae* in Japan during early, middle, and late season. Lines represent model predictions (solid: significant; dashed: non-significant). Grey areas indicate 95% CI. For abundance plots, values indicate estimated mean with 95% CI. All dots indicate individual observations. Letters indicate significant difference between means (Tukey HSD). * P < 0.05 ** P < 0.01 *** P < 0.001

Natural enemy abundance (Fig. 4e and Fig. 4m) was higher on *R. indica* growing under high water availability (GLM, water: 200 mL day^−1^, estimate = 1.98, Z = 4.21, *P* < 0.001), but lower during middle and late season (GLM, season: middle, estimate = −1.54, Z = −2.28, *P* = 0.022; season: late, estimate = −2.04, Z = −3.26, *P* = 0.001). Natural enemy abundance was lower in Taiwan compared to Japan (GLM, region: Taiwan, estimate = −1.84, Z = −3.24, *P* = 0.001), but increased drastically in middle and late season in Taiwan (GLM, season*location: middle*Taiwan, estimate = 2.51, Z = 5.04, *P* < 0.001; season*location: late*Taiwan, estimate = 4.49, Z = 8.61, *P* < 0.001).

The composition of invertebrate herbivores and natural enemies associated with *R. indica* changed as a result of water availability and season. In Taiwan, there was a clear separation of herbivores on *R. indica* growing under high and low water availability (Fig. 5a). Indicator species analysis revealed that several specialist herbivores associated significantly with *R. indica* growing under high water availability (Table 1). In contrast, generalist herbivores had a weaker association with *R. indica* growing under high water availability. Grasshoppers (Family: Catantopidae, Order: Orthoptera) were the only generalist that associated significantly with *R. indica* growing under high water availability in Japan. The only morphospecies associated with *R. indica* growing under low water availability was spider mites (Tetranychidae) in Taiwan (Table 1). In addition to water treatment, the composition of herbivores also changed across seasons in Taiwan. Both PCA (Fig. 5b) and indicator species analysis (Table 1) indicated no strong association between *R. indica* and natural enemies under low water availability. The association between natural enemies and *R. indica* were stronger in later seasons in Taiwan under high water availability (Table 1). Many of the natural enemies we observed attacked *P. rapae*. Based on field observations, several species were observed to cause high mortality of eggs and larvae of *P. rapae*, such as ants (Formicidae), predatory stink bug (*Eocanthecona concinna*), ladybugs (Coccinellidae), and spiders (Araneae). One of the surprising finding in Taiwan was a less than 1% parasitism rate by specialist parasitoid wasp, *Cotesia glomerata*, of *P. rapae* larvae on *R. indica* compared to *P. rapae* larvae in nearby cabbage fields where parasitism was regularly observed.

**Fig 5.**
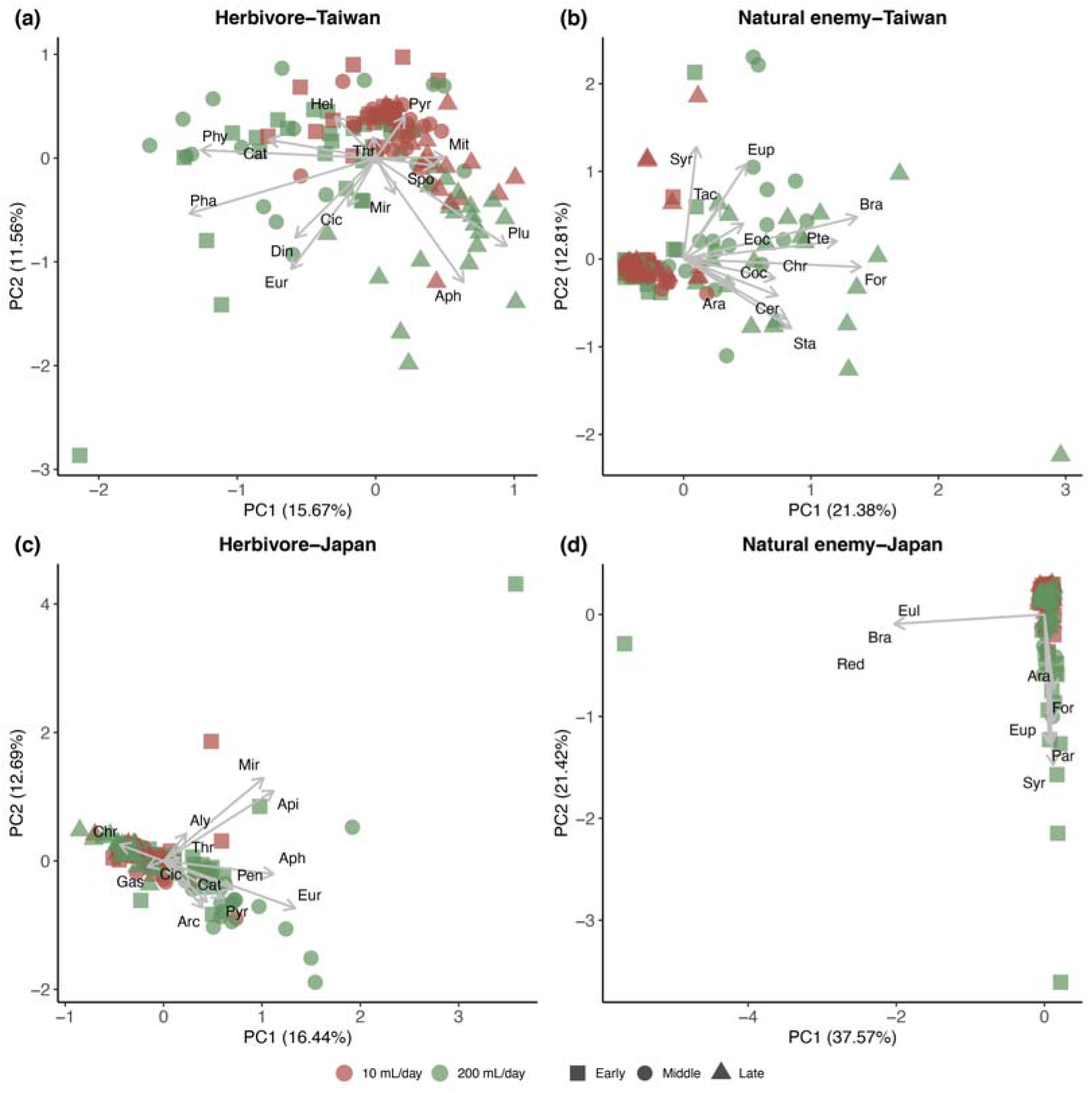
Herbivore and natural enemy compositions on *Rorippa indica* with different water status across seasons. (a) Herbivore composition in Taiwan. (b) Natural enemy composition in Taiwan. (c) Herbivore composition in Japan. (d) Natural enemy composition in Japan. For method please see Appendix 6. Abbreviation for herbivore: Aph=Aphididae, Api=Apidae, Acr=Acrididae, Aly=Alydidae, Cat=Catantopidae, Cic=Cicadellidae, Chy=Chrysomelidae, Din=Dinidoridae, Eur=*Eurydema dominulus*, Gas=Gastropoda, Hel=*Helicoverpa armigera*, Mir=Miridae, Mit=Tetranychidae, Pen=Pentatomidae, Pha=*Phaedon brassicae*, Phy=*Phyllotreta striolata*, Plu=*Plutella xylostella*, Pyr=Pyrgomorphidae, Spo=*Spodoptera litura*, Thr=Thripidae. Abbreviation for natural enemy: Ara=Araneae, Bra=Braconidae, Cer=Ceraphronidae, Chr=Chrysopidae, Coc=Coccinellidae, Eoc=*Eocanthecona concinna*, Eup=Eupelmidae, Eul=Eulophidae, For=Formicidae, Pte=Pteromalidae, Syr=Syrphidae, Tac=Tachinidae, Tro=Trombidiidae, Red=Reduviidae.

**Fig 6.**
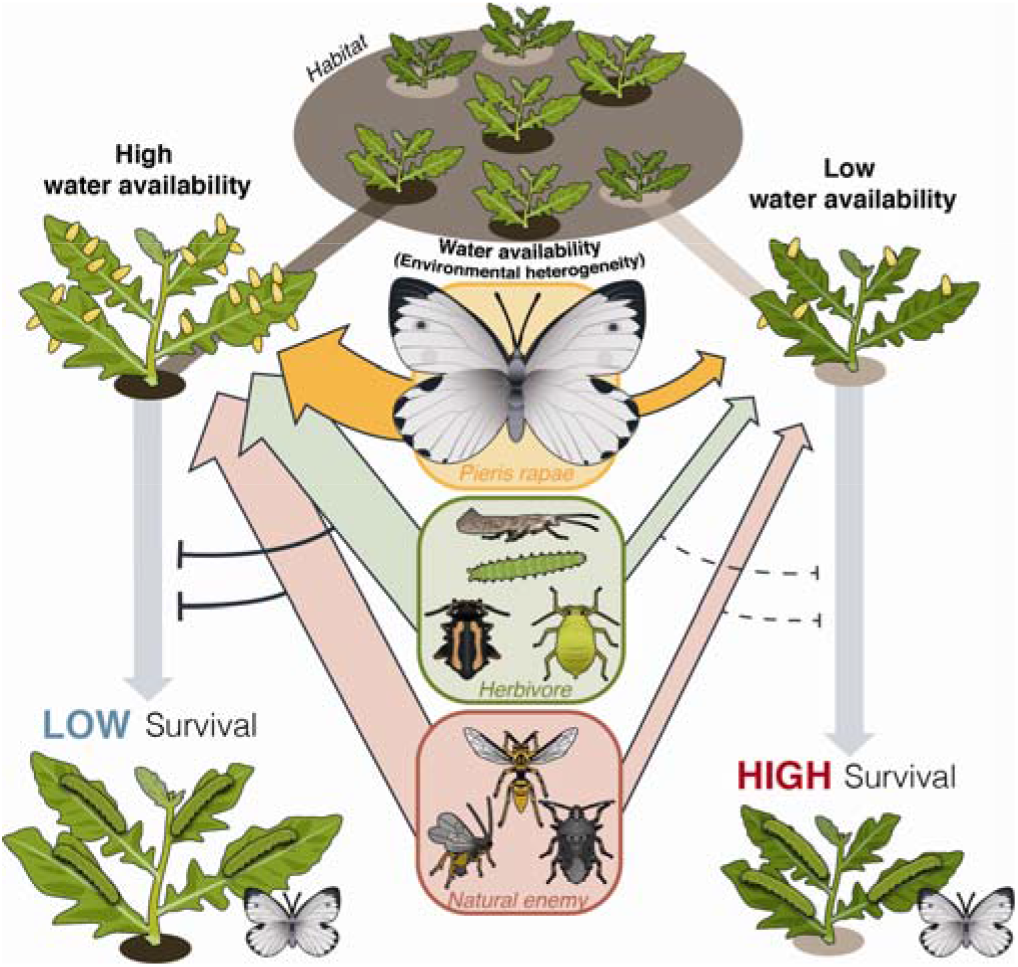
Graphical summary of relationship between water availability, plant, and insect herbivore. Female *Peiris rapae* showed ovipositional preferences toward *Rorippa indica* growing under high water availability. *R. indica* growing under low water availability were associated with lower overall abundance of arthropods. The changes in species interactions may contribute to the higher survival of *P. rapae* larvae on *R. indica* growing under low water availability.

**Table 1.**
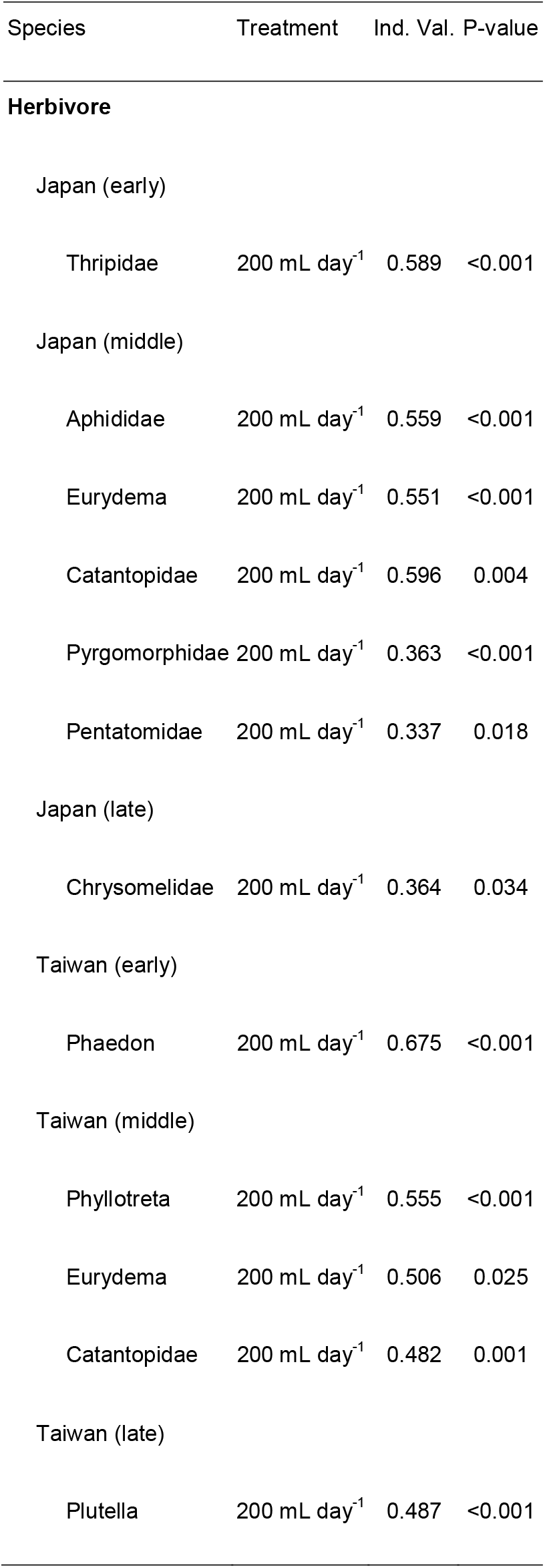
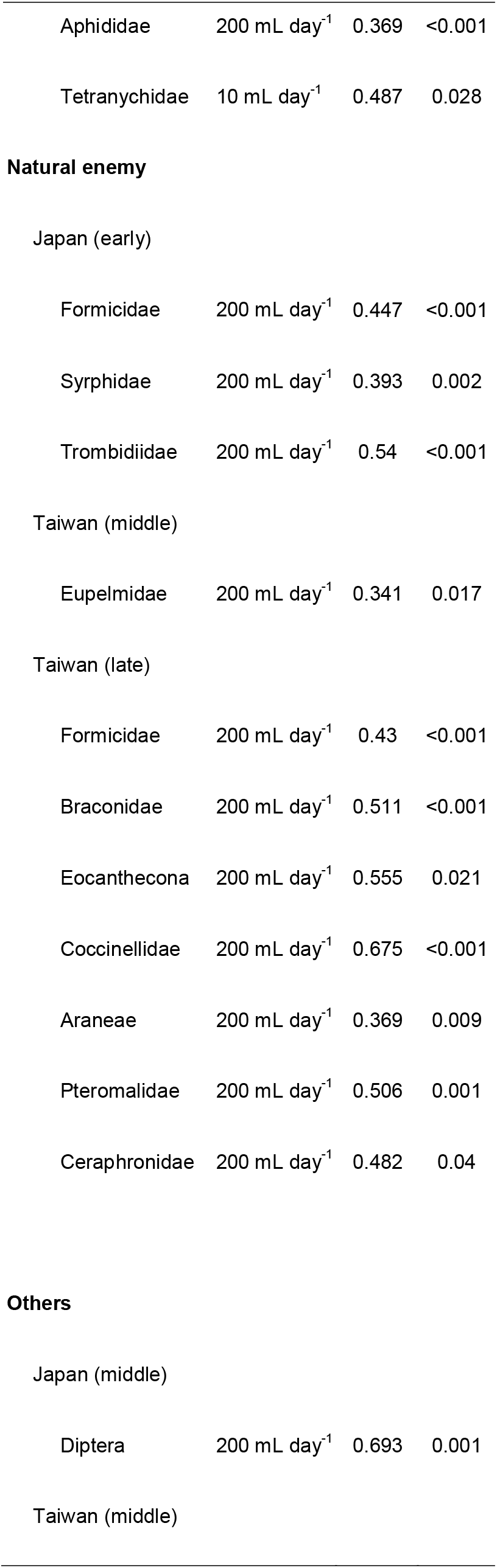
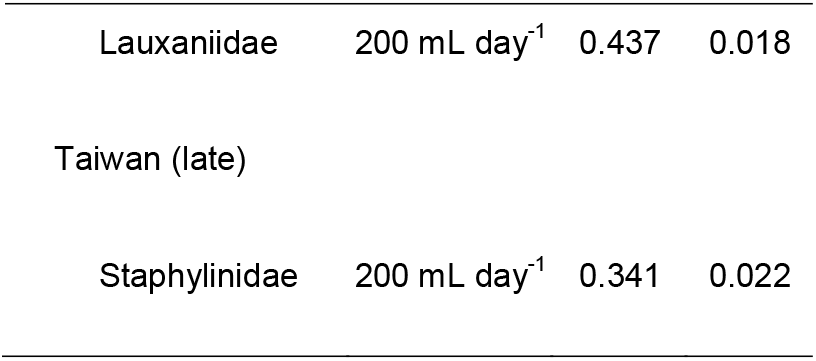
Indicator species of water availability in different season.

| Species | Treatment | Ind. Val. | P-value |
| --- | --- | --- | --- |
| **Herbivore** |  |  |  |
| Japan (early) |  |  |  |
| Thripidae | 200 mL day<sup>-1</sup> | 0.589 | <0.001 |
| Japan (middle) |  |  |  |
| Aphididae | 200 mL day<sup>-1</sup> | 0.559 | <0.001 |
| Eurydema | 200 mL day<sup>-1</sup> | 0.551 | <0.001 |
| Catantopidae | 200 mL day<sup>-1</sup> | 0.596 | 0.004 |
| Pyrgomorphidae | 200 mL day<sup>-1</sup> | 0.363 | <0.001 |
| Pentatomidae | 200 mL day<sup>-1</sup> | 0.337 | 0.018 |
| Japan (late) |  |  |  |
| Chrysomelidae | 200 mL day<sup>-1</sup> | 0.364 | 0.034 |
| Taiwan (early) |  |  |  |
| Phaedon | 200 mL day<sup>-1</sup> | 0.675 | <0.001 |
| Taiwan (middle) |  |  |  |
| Phyllotreta | 200 mL day<sup>-1</sup> | 0.555 | <0.001 |
| Eurydema | 200 mL day<sup>-1</sup> | 0.506 | 0.025 |
| Catantopidae | 200 mL day<sup>-1</sup> | 0.482 | 0.001 |
| Taiwan (late) |  |  |  |
| Plutella | 200 mL day<sup>-1</sup> | 0.487 | <0.001 |
| Aphididae | 200 mL day<sup>-1</sup> | 0.369 | <0.001 |
| Tetranychidae | 10 mL day<sup>-1</sup> | 0.487 | 0.028 |
| **Natural enemy** |  |  |  |
| Japan (early) |  |  |  |
| Formicidae | 200 mL day<sup>-1</sup> | 0.447 | <0.001 |
| Syrphidae | 200 mL day<sup>-1</sup> | 0.393 | 0.002 |
| Trombididae | 200 mL day<sup>-1</sup> | 0.54 | <0.001 |
| Taiwan (middle) |  |  |  |
| Eupelmidae | 200 mL day<sup>-1</sup> | 0.341 | 0.017 |
| Taiwan (late) |  |  |  |
| Formicidae | 200 mL day<sup>-1</sup> | 0.43 | <0.001 |
| Braconidae | 200 mL day<sup>-1</sup> | 0.511 | <0.001 |
| Eocanthecona | 200 mL day<sup>-1</sup> | 0.555 | 0.021 |
| Coccinellidae | 200 mL day<sup>-1</sup> | 0.675 | <0.001 |
| Araneae | 200 mL day<sup>-1</sup> | 0.369 | 0.009 |
| Pteromalidae | 200 mL day<sup>-1</sup> | 0.506 | 0.001 |
| Ceraphronidae | 200 mL day<sup>-1</sup> | 0.482 | 0.04 |
| **Others** |  |  |  |
| Japan (middle) |  |  |  |
| Diptera | 200 mL day<sup>-1</sup> | 0.693 | 0.001 |
| Taiwan (middle) |  |  |  |
| Lauxaniidae | 200 mL day<sup>-1</sup> | 0.437 | 0.018 |
| Taiwan (late) |  |  |  |
| Staphylinidae | 200 mL day<sup>-1</sup> | 0.341 | 0.022 |

In Japan, the separation of herbivore composition between *R. indica* growing under different water availability was less apparent (Fig. 5c). Based on indicator species analysis, most herbivores were significantly associated with *R. indica* growing under high water availability (Table 1). Season also influenced the composition of herbivore on *R. indica*. Only scarlet shieldbug (*E. dominulus*) and grasshoppers (Catantopidae) sometimes caused noticeable damage, but the level of herbivory in the field was not comparable to the level of herbivory in Taiwan. For natural enemy compositions, there were no observable differences between *R. indica* growing under different water availability (Fig. 5d). According to indicator species analysis some natural enemies were significantly associated with *R. indica* growing under high water availability in early season (Table 1). However, we did not observe any predation on *P. rapae* throughout all field experiments.

### Survival of *Pieris rapae* and arthropod community

We found significant negative associations between the abundance of herbivores and natural enemies with the survival of *P. rapae* during several seasons in both locations (Fig. 4). In Taiwan, abundance of herbivore was negatively associated with the survival of *P. rapae* during early (Fig. 4b, GLMM, estimate = −1.75, Z = −5.7, *P* < 0.001) and middle season (Fig. 4c, GLMM, estimate = −0.12, Z = −2.41, *P* = 0.016). Abundance of natural enemy was negatively associated with survival of *P. rapae* during early (Fig. 4f, GLMM, estimate = −3.64 Z = −10.11, *P* < 0.001) and late season (Fig. 4h, GLMM, estimate = −0.24, Z = −4.05, *P* < 0.001).

Most of the herbivores observed on *R. indica* showed negative associations with *R. indica* survival (Table 2). For instance, flea beetles (*P. stiolata*) that heavily attacked well-watered *R. indica* were negatively associated with survival of *P. rapae* during early season in Taiwan (GLMM, estimate = −6.50, *P* = 0.04). Two herbivores were found to be positively associated with survival of *P. rapae*, including scarlet shieldbugs (*E. dominulus*) and Cicadellidae during middle and late season in Taiwan respectively (GLMM, estimate = 0.99, p = 0.08; GLMM, estimate = 1.31, *P* < 0.001). However, the scarlet shieldbug (*E. dominulus*) had contrasting associations with the survival of *P. rapae* in different month (negative association during late season, GLMM, estimate = −0.95, *P* < 0.001), likely due to the difference in abundance between seasons (Individual plant^−1^: Middle = 0.136; Late = 0.275). The analysis of morphospecies in Japan revealed some unexpected associations; for example, Lauxaniidae, a parasitic mite of aphids, and Syrphidae (during early season, GLMM, estimate = −0.95, *P* = 0.021; estimate = 0.50, p = 0.001; estimate = −0.31, *P* = 0.041).

**Table 2.**
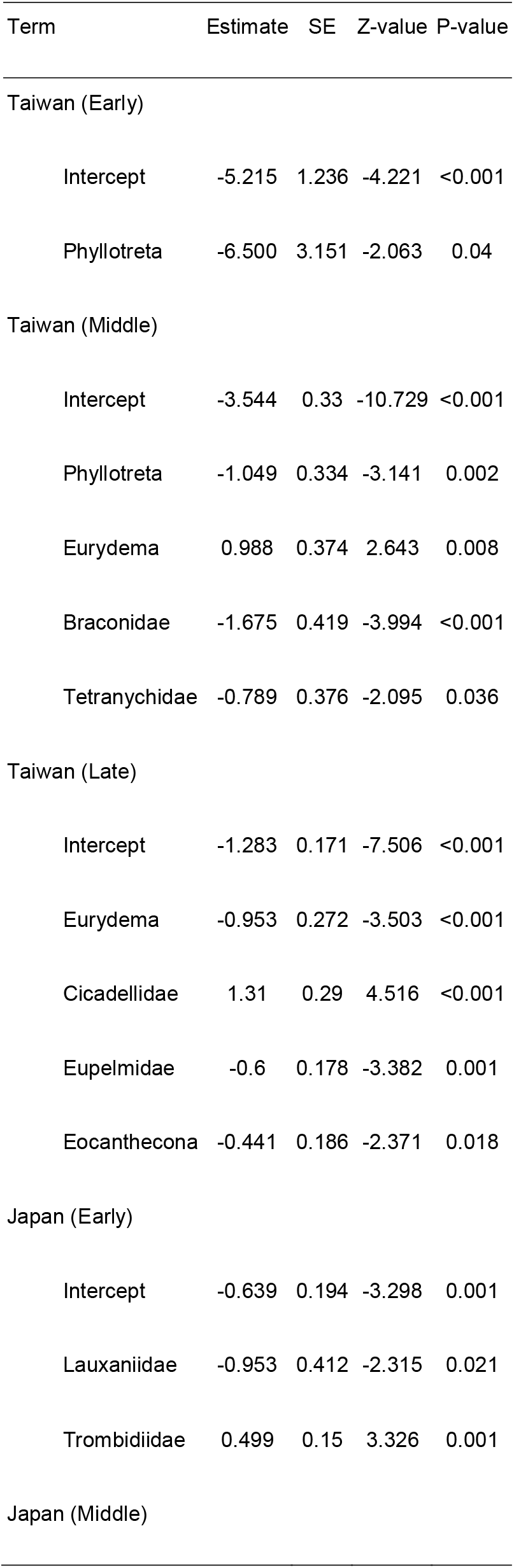
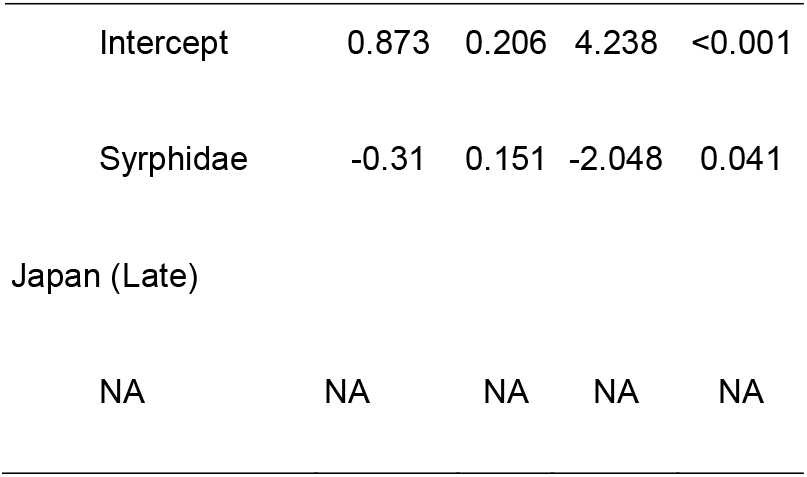
Model summaries of associations between survival of Pieris rapae and morphospecies.

| Term | Estimate | SE | Z-value | P-value |
| --- | --- | --- | --- | --- |
| Taiwan (Early) |  |  |  |  |
| Intercept | -5.215 | 1.236 | -4.221 | <0.001 |
| Phyllotreta | -6.500 | 3.151 | -2.063 | 0.04 |
| Taiwan (Middle) |  |  |  |  |
| Intercept | -3.544 | 0.33 | -10.729 | <0.001 |
| Phyllotreta | -1.049 | 0.334 | -3.141 | 0.002 |
| Eurydema | 0.988 | 0.374 | 2.643 | 0.008 |
| Braconidae | -1.675 | 0.419 | -3.994 | <0.001 |
| Tetranychidae | -0.789 | 0.376 | -2.095 | 0.036 |
| Taiwan (Late) |  |  |  |  |
| Intercept | -1.283 | 0.171 | -7.506 | <0.001 |
| Eurydema | -0.953 | 0.272 | -3.503 | <0.001 |
| Cicadellidae | 1.31 | 0.29 | 4.516 | <0.001 |
| Eupelmidae | -0.6 | 0.178 | -3.382 | 0.001 |
| Eocanthecona | -0.441 | 0.186 | -2.371 | 0.018 |
| Japan (Early) |  |  |  |  |
| Intercept | -0.639 | 0.194 | -3.298 | 0.001 |
| Lauxaniidae | -0.953 | 0.412 | -2.315 | 0.021 |
| Trombididae | 0.499 | 0.15 | 3.326 | 0.001 |
| Japan (Middle) |  |  |  |  |
| Intercept | 0.873 | 0.206 | 4.238 | <0.001 |
| Syrphidae | -0.31 | 0.151 | -2.048 | 0.041 |
| Japan (Late) |  |  |  |  |
| NA | NA | NA | NA | NA |

## Discussion

As spatial and temporal variation in plant access to water increases in the future (National Research Council 2011), understanding the impacts of water availability on plants and the associated heterotrophs are important steps to identify complex effects of water availability on community interactions. By evaluating interactions across multitrophic levels, we show that plant water availability differentially impacts specific herbivores and has overall effects on arthropod community composition. Low water availability reduces overall arthropod abundance associated with the plants, and this negative impact is linked to decrease in plant quality indicated by reduction in caterpillar performances. These findings are similar to the common observation that low water availability affects insect herbivores on herbaceous plant negatively (Waring and Cobb 1992). Notably, we observed that a specialist herbivore was capable of coping with fluctuations in host plant quality driven by water availability. The performance of *P. rapae* from Taiwan and Japan revealed a potential case of local adaptation, where *P. rapae* in Taiwan and Japan performed equally well on sympatric *R. indica* of different water status. The ability for *P. rapae* to cope with variation in host quality, suggests the role of host variations in shaping the evolution of physiological capacity of *P. rapae*. Similar local specialization caused by regional effects and host plants was also documented in other butterfly, such as *Melitaea cinxia* (Kuussaari et al. 2000).

Even with the physiological capacity to overcome quality variations of plant, *P. rapae* still displayed preference toward *R. indica* growing under high water availability. As predicted by preference-performance hypothesis (PPH), oviposition of females should match the performance of their offspring (Gripenberg et al. 2010; Thompson 1988). Based on the observed decrease in oviposition on *R. indica* growing under low water availability in both populations, the preference-performance relationship predicted by PPH was not observed. It has been shown that the preference of females could be blurred by temporal variation in resource quality (Cronin et al. 2001; Gripenberg et al. 2007), and thus failure to evolve a proper mechanism to perceive these changes in host quality for their offspring. Alternatively, this behavior could also be a passive response of female *P. rapae* to changes in attractiveness (due to size or chemical cues) of plants under low water availability. For example, changes in volatile emission patterns are known to be important in determining the attractiveness of plants to insects (Orre et al. 2010; Pichersky and Gershenzon 2002), and water availability affects volatile emissions (Copolovici et al. 2014; Scott et al. 2018). It is also likely that food source stability (Garssen et al. 2014), or preference for food with higher water content have led to the evolution of preference toward plants under high water availability.

Given the capacity of *P. rapae* to utilize host plants of variable water status, the reduction in other herbivores and natural enemies on *R. indica* growing under low water availability emerged as an important factor that increases the survival of *P. rapae*. Although we did not explicitly evaluate the level of predation and competition, the drastic changes in abundance of herbivores and natural enemies provides a strong basis in shifting in competition and predation. Several morphospecies that either consistently observed to prey on *P. rapae* or cause large amounts of damage to *R. indica* in Taiwan, were linked to observable change in *P. rapae* abundance on *R. indica*. In addition, the overall low abundance of both herbivores and natural enemies, and species that directly interact with *P. rapae* in Japan, is a likely reason of the lack of overall association between abundance of arthropods and survival of *P. rapae* on *R. indica*. This result further supports our conclusion that multitrophic interaction is an important factor contributing to the benefit of low water availability to *P. rapae*.

A surprising finding that appeared was the positive associations between the survival of *P. rapae* with two piercing-sucking herbivores in Taiwan. Although this observation contradicted our overall conclusion that stronger competition reduced the survival of *P. rapae* on *R. indica* growing under high water availability, it reveals potential divergent defense responses elicited by herbivores in different feeding guilds. For example, plant defenses induced by piercing-sucking hemipterans are often salicylic acid-dependent, in contrast to being jasmonic acid-dependent in chewing herbivores. The antagonistic relationship between these defenses has been frequently shown to benefit chewing herbivores (De Vos et al. 2005; Kroes et al. 2015; Li et al. 2014; Pieterse et al. 2012; Soler et al. 2012). We speculate that the positive associations between the survival of *P. rapae* and two piercing-sucking herbivores were likely mediated by a similar mechanism described above. It has also been shown that the positive impacts of piercing-sucking herbivores took place only under low density (Kroes et al. 2015), which explains the switch in association between survival of *P. rapae* and *E. dominulus* from positive to negative as the density of *E. dominulus* doubled. Further exploration into mechanisms and associated tradeoffs in these interactions are needed.

There are some caveats to our study. First, while we generated plants of different water status morphologically comparable to field individuals, these methods for quantifying water status are mostly indirect measures of water stress rather than detailed physiological characterizations of plant stress (Jones 2007). Although they are valuable in detecting plant responses to water limitation, they are not useful for studies investigating changes in physiological/molecular processes associated with droughts (Jones 2007). Second, although using detached leaves in caterpillar RGR bioassay has been shown to reflect chemical changes in leaves (Chung et al. 2013; Wang et al. 2017), we cannot rule out that detaching leaf itself resulted in alterations in leaf quality. While this method enables us to state there are changes in the host plants, it may not fully reflect the herbivores performance under natural feeding conditions. Third, since we only recorded the abundance of herbivore and natural enemies once in each field experiment, it is possible that arthropod abundance and species richness are underestimated. Finally, the time period of the field experiment was shorter than the larval stage of *P. rapae*. As a result, we do not know whether *R. indica* can support the number of larvae observed in field. However, the likelihood of complete defoliation seems low due to the ability of plants to regrow and presence of natural enemies under high water availability, and the low quality of plants to other herbivores under low water availability. The majority of the plants still maintained approximately 50-70% of its biomass after the 10 days field experiment.

The observation that certain herbivores (i.e. *P. rapae*) were unaffected by the reduction in host quality suggests that addressing focal herbivores without incorporating the natural insect community can alter interpretations of herbivore success. It can be misleading to focus on few selected insect species and groups of defense responses because changes in plants under low water availability might have differential impacts to community members. Some insect herbivores might have better ability to cope with changes in plant quality than others, while others may be negatively affected by it. Future studies should aim to include more focal species and aspects of plant-insect interactions in order to capture a broader scope of how insect and plant communities respond to environmental changes.

In summary, the present study provides insights to the ongoing discussion on how water availability influences insect herbivores (Jamieson et al. 2012; Mattson and Haack 1987; White 2009). By studying impact of water availability under the context of arthropod community in addition to the direct interaction between plants and insect herbivores, we revealed a potential factor in which plants growing under low water availability can be beneficial to some insects but detrimental to others. The contradicting impacts of water availability on different arthropod species highlight the complexity and context dependency of these interactions. The findings also beg the questions of how variation caused by environmental heterogeneity, including heterogeneity in water availability and other factors, affects the ecology and evolution of interactions between plants and insect herbivores.

## Acknowledgement

We thank the International Agriculture and Development Graduate Program (Pennsylvania State University) for providing monetary support and feedback on experimental design. We thank Chi-Shun Chang, Yi-Zhang Wang, Han-Rong Li, Kazumu Kuramitsu, and Kai Han for collecting data. We would like to express special thanks to Dr. Yooichi Kainoh for arranging space and logistics of experiment in Japan, Dr. Chin-Wen Tan and Wei-Ting Chen for helpful information on the experiments in Taiwan, and Dr. Charles Mason and Jagdeep Singh Sidhu for feedbacks on manuscript.

## Data availability statement

Data in this manuscript will be deposited and made available in the Dryad Digital Repository.

## Declaration of authorship

PAL, KN, CKH, WPC, and GWF conceived the ideas, designed methodology. PAL and JAO analyzed the data. PAL, CML, and CHS collected the data. PAL, CML, JAO, KN, CKH, WPC, and GWF wrote the manuscript.

## Notes

### Competing Interest Statement

The authors have declared no competing interest.

