## Supplemental information for "Changes in arthropod community but not plant quality benefit a specialist herbivore on plant under reduced water availability"

### Supporting information

#### Appendix 1 figure

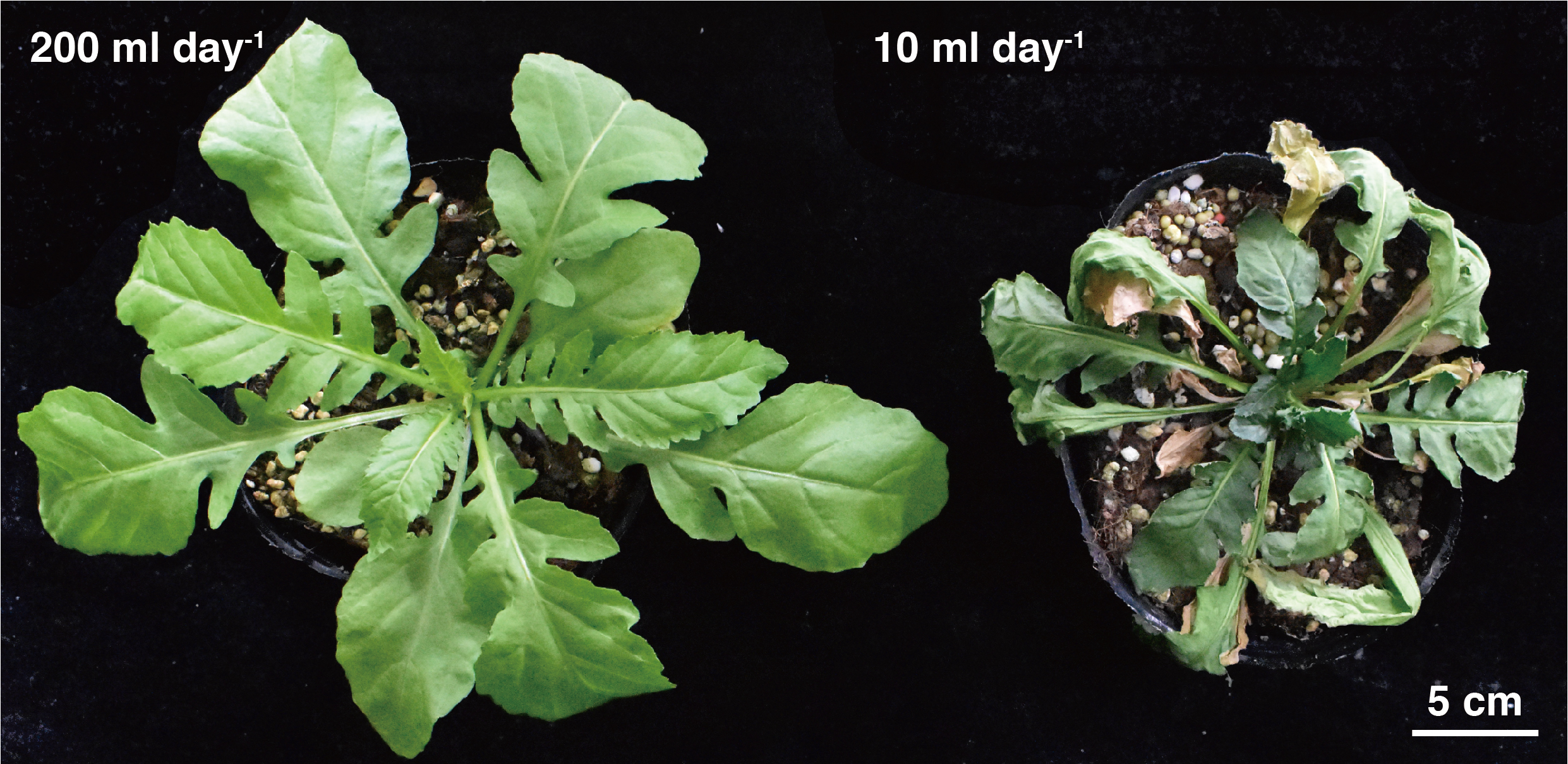

Appendix 1 *Rorippa indica* growing under different water availability in field. Picture was taken on a sunny day at 12 p.m. in Taiwan. Low water availability (on the right) phenotype were observed at 12, 2, and 4 p.m. to prevent tissue damage. If the meristem showed wilting symptom (as in the photo) 10 ml of water was added at the time of observation. The phenotype was maintained as consistent as possible throughout the field experiment.

#### Appendix 2 figure

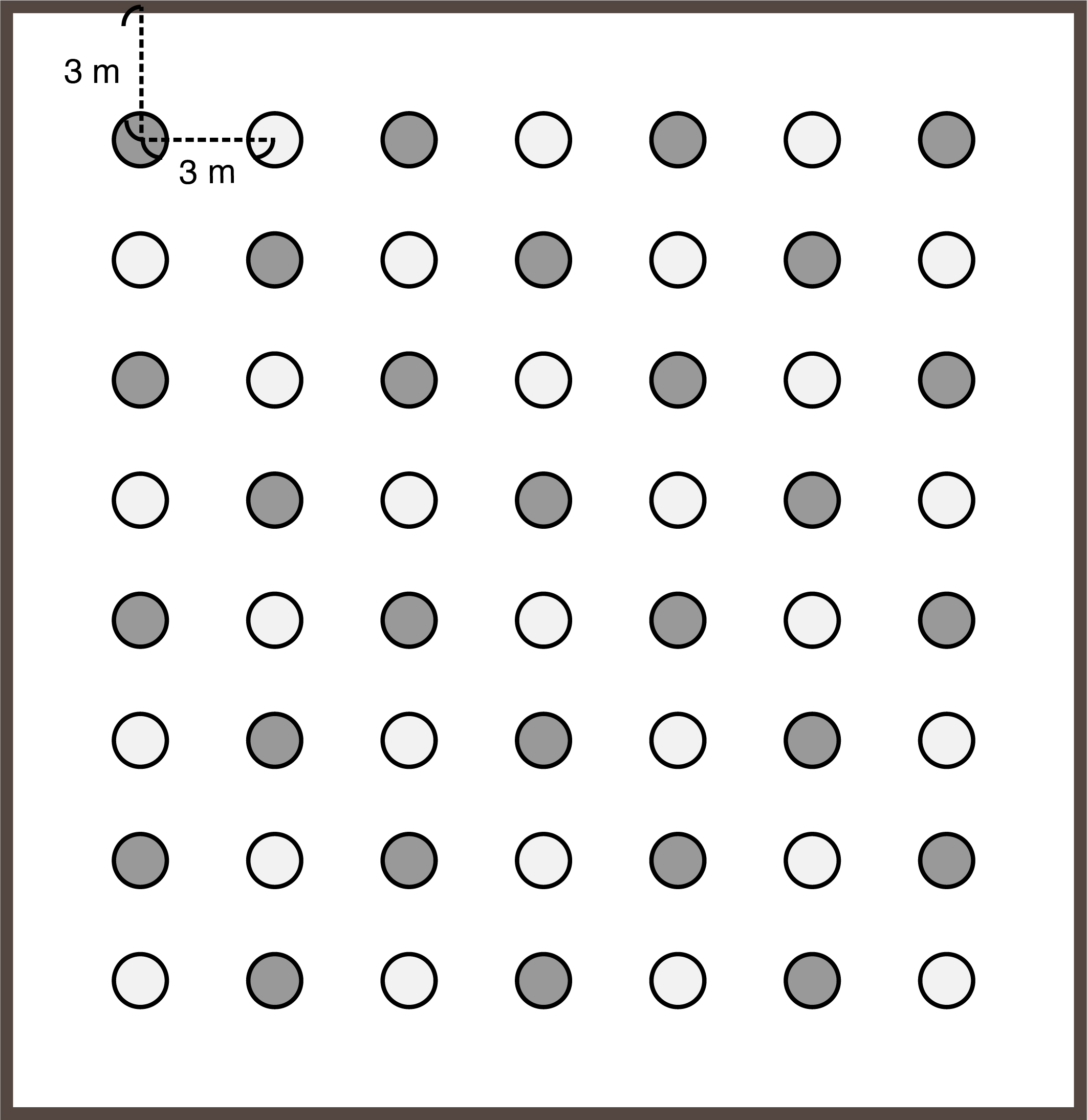

Appendix 2 Layout of experimental field. Dark grey circles indicate low water availability; light grey circles indicate high water availability*.* Each plant was 3 meters apart, and 3 meters away from the border of field. During field observations, 2-3 people were randomly assigned to rows, each person starts the observation at different directions (left to right or vice versa) to minimize the potential variations caused by disturbance of human activity. We also tried to lower the level of disturbance during the observations to minimize artifacts.

#### Appendix 3 table

Appendix 3 list of morphospecies observed in field.

| Category | Location | Morphospecies |
| --- | --- | --- |
| Herbivore | Japan | Aphididae |
|  |  | *Eurydema dominulus* |
|  |  | Thripidae |
|  |  | Chrysomelidae |
|  |  | Unknown caterpillar |
|  |  | Cicadellidae |
|  |  | Acrididae |
|  |  | Catantopidae |
|  |  | Pyrgomorphidae |
|  |  | Pentatomidae |
|  |  | Miridae |
|  |  | Apidae |
|  |  | Gastropoda |
|  | Taiwan | *Phyllotreta striolata* |
|  |  | *Phaedon brassicae* |
|  |  | *Plutella xylostella* |
|  |  | Aphididae |
|  |  | *Eurydema dominulus* |
|  |  | Thripidae |
|  |  | *Helicoverpa armigera* |
|  |  | *Spodoptera litura* |
|  |  | Cicadellidae |
|  |  | Catantopidae |
|  |  | Pyrgomorphidae |
|  |  | Miridae |
|  |  | Dinidoridae |
|  |  | Tetranychidae |
| Natural enemy | Japan | Formicidae |
|  |  | Braconidae |
|  |  | Trombidiidae |
|  |  | Eupelmidae |
|  |  | Syrphidae |
|  |  | Reduviidae |
|  |  | Eulophidae |
|  |  | Araneae |
|  | Taiwan | Formicidae |
|  |  | Braconidae |
|  |  | Ceraphronidae |
|  |  | Chrysopidae |
|  |  | Eupelmidae |
|  |  | Syrphidae |
|  |  | *Eocanthecona concinna* |
|  |  | Tachinidae |
|  |  | Coccinellidae |
|  |  | Araneae |
|  |  | Pteromalidae |
| Others | Japan | Lauxaniidae |
|  |  | Unknown Diptera |
|  |  | Apidae |
|  | Taiwan | Lauxaniidae |
|  |  | Unknown Diptera |
|  |  | Collembolla |
|  |  | Chironomidae |
|  |  | Staphylinidae |
